# Neurodegenerative and functional signatures of the cerebellar cortex in m.3243A>G patients

**DOI:** 10.1101/2021.04.30.442091

**Authors:** Roy AM. Haast, Irenaeus FM. De Coo, Dimo Ivanov, Ali R Khan, Jacobus FA. Jansen, Hubert JM. Smeets, Kâmil Uludağ

## Abstract

Mutations of the mitochondrial DNA are an important cause of inherited diseases that can severely affect the tissue’s homeostasis and integrity. The m.3243A>G mutation is the most commonly observed across mitochondrial disorders and is linked to multisystemic complications, including cognitive deficits. In line with in vitro experiments demonstrating the m.3243A>G’s negative impact on neuronal energy production and integrity, m.3243A>G patients show cerebral gray matter tissue changes. However, its impact on the most neuron-dense, and therefore energy-consuming brain structure – the cerebellum – remains elusive.

In this work, we used high resolution structural and functional data acquired using 7 Tesla MRI to characterize the neurodegenerative and functional signatures of the cerebellar cortex in m.3243A>G patients. Our results reveal altered tissue integrity within distinct clusters across the cerebellar cortex, apparent by their significantly reduced volume and longitudinal relaxation rate compared to healthy controls, indicating macroscopic atrophy and microstructural pathology. Spatial characterization reveals that these changes occur especially in regions related to the frontoparietal brain network that is involved in information processing and selective attention. In addition, based on resting-state fMRI data, these clusters exhibit reduced functional connectivity to frontal and parietal cortical regions, especially in patients characterized by (i) a severe disease phenotype and (ii) reduced information processing speed and attention control.

Combined with our previous work, these results provide insight into the neuropathological changes and a solid base to guide longitudinal studies aimed to track disease progression.

## Introduction

Among the many mitochondrial mutations reported,^1^ the adenine (A) to guanine (G) transition at base pair 3243 within the *MT-TL1* gene encoding tRNALeu^(UUR)^, better known as the m.3243A>G mutation, has been commonly observed across the spectrum of mitochondrial disorders.^2, 3^ Its clinical expression varies strongly, ranging from patients that are non-symptomatic to patients suffering from episodes of severe stroke-like symptoms.^4^ The most prominent symptoms are hearing loss (48%), gastro-intestinal symptoms (42%), decreased vision (42%), exercise intolerance (38%), glucose intolerance (37%), gait instability (36%), cerebellar ataxia (35%), myopathy (34%), cognition impairment (32%) and ptosis (32%).^5^ In symptomatic patients, the collection of symptoms are often incorrectly referred to as the ‘mitochondrial encephalopathy lactic acidosis and stroke-like episodes (MELAS)’,^6^ as stroke-like episodes are only present in 4% of the patients,^7^ or ‘maternally inherited diabetes and deafness’^8^ syndrome. Despite its relatively high prevalence compared to other mitochondrial mutations, descriptions of neuroradiological changes in m.3243A>G patients are predominantly based on single-case neuroimaging studies and only a limited number of studies have focused on larger cohorts.^9–15^

We have previously reported on the structural changes across the cerebral cortex and subcortical nuclei in a relatively large cohort of twenty-two m.3243A>G patients using high resolution, quantitative 7 Tesla (7T) MRI data.^16^ We found significant volume, microstructural and perfusion differences in the brains of patients compared to healthy controls and showed that the magnitude of cerebral gray matter (GM) changes with the percentage affected mitochondria per cell (i.e., ‘mutation load’ or ‘heteroplasmy rate’) and disease severity. Here, specific cortical regions, linked to attentional control (e.g., middle frontal gyrus), the sensorimotor network (e.g., banks of central sulcus) and the default mode network (e.g., precuneus) were shown more prone for affected tissue integrity.

Despite the sparse, but growing knowledge about the impact on the cerebral cortex, the neuroradiological correlates of the cerebellum of the m3243A>G mutation continue to remain understudied. Earlier *ex vivo* work has revealed a wide range of neuropathological findings in cerebellar tissue taken from m.3243A>G patients.^17^ Given the crucial role of mitochondria energy production in neuronal survival^18^ a detailed*, in vivo* characterization of cerebellar tissue changes may provide complementary insight in the neuropathological expression of the m.3243A>G mutation and its effect on overall brain’s functioning. The cerebellum features the most strongly convoluted GM across the entire human brain with densely packed neurons that together account for 78 % of the brain’s entire surface area.^19^ Traditionally, it is linked to sensorimotor control, ensuring coordinated and timed movements,^20^ but its prominence across a broader range of cognitive processes has recently been confirmed through the characterization of its functional topography.^21^ Here, distinct regions within the cerebellar GM are involved in a diverse set of motor, cognitive, and social and affective tasks and confirm earlier initial findings.^22–24^ As such, impaired cerebellar connectivity due to disease may have profound implications for the integrity of motor and non-motor brain networks.^25^

In this study, we extend our initial cerebral work with previously unexplored high resolution functional 7T MRI data to characterize (i) macroscopic and microstructural changes in the cerebellum of m.3243A>G patients and explore their (ii) spatial correspondence with the cerebellar’s anatomical and functional parcellation, (iii) effect on functional cerebello–cortical connectivity and (iv) correlation with disease severity and cognitive outcome measures. The presented results demonstrate a first and unique description of the neurodegenerative and functional signatures of the cerebellum related to the m.3243A>G mutation.

## Materials and methods

### Subject recruitment

Twenty-two m.3243A>G patients and fifteen healthy controls were included in this study after providing written informed consent in accordance with the Declaration of Helsinki. The experimental procedures were approved by the ethics review board of the MUMC+ in Maastricht, The Netherlands. Participants were matched based on age, gender and education (see Table 1). A more detailed description of the in- and exclusion criteria, as well as patient characteristics can be reviewed in an earlier manuscript.^16^ Most importantly, disease severity scores were obtained (i) by an experienced clinician (I.F.M.d.C) using the Newcastle Mitochondrial Disease Adult Scale (NMDAS, see Supplementary Table 1)^26^ and (ii) m.3243A>G mutation loads in urine epithelial cells (UECs) and blood, corrected for age and sex, respectively.^27^ Subjects in the acute phase or with a history of SLEs based on the Barthel (i.e. ADL-independent) and NMDAS (<30) criteria) were excluded resulting in a in a spectrum with less severe phenotypes and without a diagnosis of a cerebellar motor deficit. Cognitive performance scores were collected to correlate with MRI-based findings. This included the letter-digit-substitution task (LDST) to test information processing speed, the Stroop colour-word task to test attention and the visual 15-Words Learning Task (15-WLT) to test memory, recall and recognition.^28–30^ Raw test scores were Z-scored based on the average control scores for each cognitive task (see Table 1). None of the patients reported subjective cognitive difficulties.

**Table 1.**
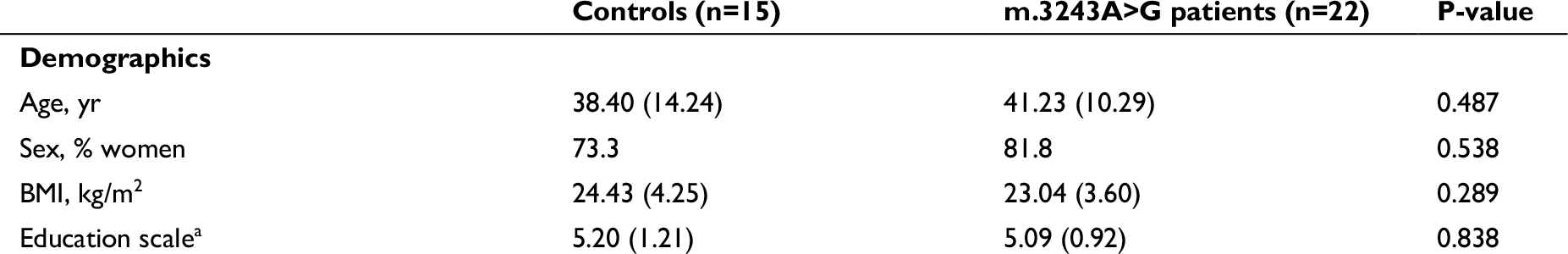
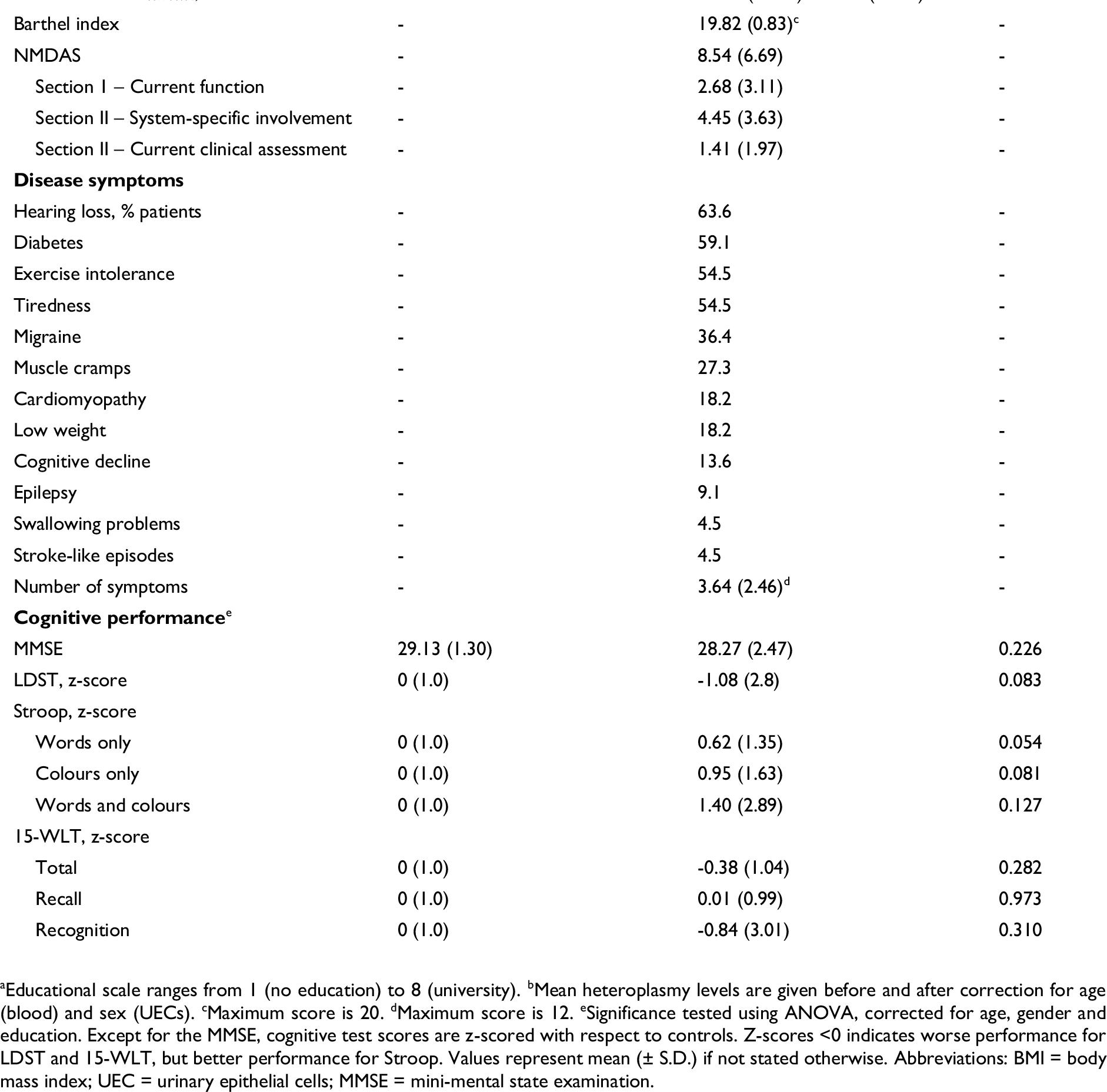
Study population demographics.

|  | Controls (n=15) | m.3243A>G patients (n=22) | P-value |
| --- | --- | --- | --- |
| **Demographics** |  |  |  |
| Age, yr | 38.40 (14.24) | 41.23 (10.29) | 0.487 |
| Sex, % women | 73.3 | 81.8 | 0.538 |
| BMI, kg/m<sup>2</sup> | 24.43 (4.25) | 23.04 (3.60) | 0.289 |
| Education scale<sup>a</sup> | 5.20 (1.21) | 5.09 (0.92) | 0.838 |
| Mutation load<sup>b</sup> |  |  |  |
| UECs / UECs<sub>corrected</sub>, % | 0 | 53.14 (26.09) / 59.77 (26.45) | - |
| Blood / Blood<sub>corrected</sub>, % | 0 | 20.23 (11.40) / 63.11 (27.38) | - |
| Barthel index | - | 19.82 (0.83)<sup>c</sup> | - |
| NMDAS | - | 8.54 (6.69) | - |
| Section I – Current function | - | 2.68 (3.11) | - |
| Section II – System-specific involvement | - | 4.45 (3.63) | - |
| Section II – Current clinical assessment | - | 1.41 (1.97) | - |
| Disease symptoms |  |  |  |
| Hearing loss, % patients | - | 63.6 | - |
| Diabetes | - | 59.1 | - |
| Exercise intolerance | - | 54.5 | - |
| Tiredness | - | 54.5 | - |
| Migraine | - | 36.4 | - |
| Muscle cramps | - | 27.3 | - |
| Cardiomyopathy | - | 18.2 | - |
| Low weight | - | 18.2 | - |
| Cognitive decline | - | 13.6 | - |
| Epilepsy | - | 9.1 | - |
| Swallowing problems | - | 4.5 | - |
| Stroke-like episodes | - | 4.5 | - |
| Number of symptoms | - | 3.64 (2.46)<sup>d</sup> | - |
| Cognitive performance<sup>e</sup> |  |  |  |
| MMSE | 29.13 (1.30) | 28.27 (2.47) | 0.226 |
| LDST, z-score | 0 (1.0) | -1.08 (2.8) | 0.083 |
| Stroop, z-score |  |  |  |
| Words only | 0 (1.0) | 0.62 (1.35) | 0.054 |
| Colours only | 0 (1.0) | 0.95 (1.63) | 0.081 |
| Words and colours | 0 (1.0) | 1.40 (2.89) | 0.127 |
| 15-WLT, z-score |  |  |  |
| Total | 0 (1.0) | -0.38 (1.04) | 0.282 |
| Recall | 0 (1.0) | 0.01 (0.99) | 0.973 |
| Recognition | 0 (1.0) | -0.84 (3.01) | 0.310 |

<sup>a</sup>Educational scale ranges from 1 (no education) to 8 (university). <sup>b</sup>Mean heteroplasmy levels are given before and after correction for age (blood) and sex (UECs). <sup>c</sup>Maximum score is 20. <sup>d</sup>Maximum score is 12. <sup>e</sup>Significance tested using ANOVA, corrected for age, gender and education. Except for the MMSE, cognitive test scores are z-scored with respect to controls. Z-scores <0 indicates worse performance for LDST and 15-WLT, but better performance for Stroop. Values represent mean ( ± S.D.) if not stated otherwise. Abbreviations: BMI = body mass index; UEC = urinary epithelial cells; MMSE = mini-mental state examination.

### MRI acquisition

MRI data were acquired using a whole-body 7T magnet (Siemens Healthineers, Erlangen, Germany) equipped with a 32-channel phased-array head coil (Nova Medical, Wilmington, MA, USA). High resolution (0.7 mm isotropic nominal voxel size) whole-brain quantitative R_1_ and B1^+^ maps (2 mm isotropic nominal voxel size) were obtained using the 3D MP2RAGE^31^ and 3D Sa2RAGE^32^ sequences, respectively. R_1_ is an intrinsic property (i.e., longitudinal relaxation rate) of brain tissue that can be quantified using MRI and relates to tissue integrity (e.g., it decreases with demyelination).^33^ In addition to the anatomical scans, whole-brain resting-state functional MRI (rs-fMRI) data with an 1.4 mm isotropic nominal voxel size were acquired using a 2D Multi-Band Echo Planar Imaging (2D MB-EPI) sequence to probe functional connectivity between cerebellar and cortical areas. Five additional volumes were acquired with reverse phase encoding to correct the functional data for EPI readout-related geometrical distortions. See Supplementary Table 2 for the relevant sequence parameters. Dielectric pads water were placed proximal to the temporal lobe and cerebellar areas to improve image homogeneity across the brain.^34^

### MRI data analysis

In brief, anatomical data were used to extract cerebral and cerebellar cortical GM segmentations (and surfaces) for voxel-based morphometry (VBM), while the rs-fMRI data were preprocessed to assess cerebello–cortical functional connectivity.

### Anatomical data preprocessing

MP2RAGE anatomical data were preprocessed as described previously, including the removal of non-brain tissue,^35^ correction for image inhomogeneities^36, 37^ and cortical surface reconstruction and parcellation using FreeSurfer (v6.0).^38^ Native-resolution surface meshes (∼164k vertices) were downsampled to the ‘fsLR’ surface space (∼32k vertices) using the instructions and transforms provided by the Human Connectome Project (https://github.com/Washington-University/HCPpipelines).^39^

### Voxel-based morphometry workflow

Cerebellar neuroradiological changes in m.3243A>G patients were studied using the SUIT (v3.2, www.diedrichsenlab.org/imaging/suit.html) and VBM toolboxes in SPM12 through normalization to a spatially unbiased template of the cerebellum.^40, 41^ The cerebellar GM, WM and CSF masks were obtained using the cerebellar segmentation (CERES) tool,^42^ to match the previously used labels.^16^ The sum of cerebellar GM and WM maps served as the cerebellar isolation mask and were individually checked and manually corrected using ITK-SNAP (v3.6.0) to exclude non-cerebellar tissue.^16, 43^ Diffeomorphic anatomical registration (DARTEL)^44^ was employed to normalize the individual subject’s cerebellum GM and WM masks to the corresponding probability maps of the SUIT atlas. A detailed description of the underlying workflow can be found in Diedrichsen et al.^45^. The resulting deformation fields were then used to deform the tissue probability and R_1_ maps from each individual participant. Finally, transformed GM and WM probability images were multiplied by the relative voxel volumes (i.e., the Jacobian determinants of the deformation field) to correct for volume changes during the spatial normalization step^46^ and all output was spatially smoothed with a kernel of 4 mm^3^. As a result, differences in intensities marked approximate GM or WM densities (and thus served as a proxy for tissue volume changes), and R_1_ for each voxel. These could then be used to directly examine differences between patients and controls (see section on statistical analyses for further details).

### Resting-state fMRI analysis

Preprocessing of the rs-fMRI EPI volumes included slice-timing correction (using AFNI’s ‘3dTshift’),^47^ followed by estimation of (i) volume-specific motion parameter matrices (FSL’s ‘mcflirt’);^48^ (ii) gradient non-linearity (Human Connectome Project’s ‘gradient_unwarp.py’); (iii) EPI readout-related (using the opposite phase encoding images and FSL’s ‘topup’) distortions maps;^49^ and (iv) the transformation to a 1.4 mm^3^ MNI template space. To achieve the latter, first, a linear coregistration between the subject’s mean rs-fMRI EPI volume and the subject’s native skull stripped anatomical volume (i.e., EPI-to-anatomical registration, and its inverse) was calculated using FreeSurfer’s boundary-based registration implementation (‘bbregister’).^50^ This was followed by computing the subject’s native anatomical-to-MNI non-linear transformation warp (and its inverse) using FSL’s ‘fnirt’.^51^ Finally, each slice-timing-corrected rs-fMRI EPI volume was resampled and resliced into the MNI template space using a one-step procedure that included: (i) motion correction, (ii) gradient non-linearity, (iii) readout distortion and (iv) the MNI-space transformation.

Within the CONN functional connectivity toolbox (https://web.conn-toolbox.org/),52 resampled rs-fMRI data were then denoised using aCompCor (WM and CSF ROIs, five components each)^53^, scrubbing (number of identified invalid scans), motion regression (12 regressors: six motion parameters + six first-order temporal derivatives), temporal band-pass filtering (0.08 – 0.8 Hz), detrended and demeaned. In parallel, left and right hemisphere cortical (i.e., fsLR) surfaces were transformed to MNI space using the obtained inverse EPI-to-anatomical transformation matrices and Connectome Workbench’s ‘surface-apply-warpfield’ command for projection of the denoised data onto the surface.^54^

For the first-level (i.e., region of interest [ROI]-to-ROI) analyses, cerebello–cortical connectivity (i.e., correlation) matrices were computed for each subject. Here, ROIs included the cerebellar ROIs based on the VBM results of the anatomical data as well as predefined cortical ROIs based on the Schaefer (*n_regions_*=100) atlas.^55^ The Schaefer atlas exploits local gradients in resting-state functional connectivity, while maximizing similarity of rs-fMRI time courses within a parcel. It additionally allows stratification of results based on seven large-scale networks: default-mode (DMN), frontoparietal (FPN), dorsal attention (DAN), ventral attention (VAN), somatosensory (SMN), limbic and visual networks.^56^

### Statistical analyses

Group and disease severity effects were explored using the outputs from the volumetric, VBM and rs-fMRI workflows above and the statistical models implemented in the statsmodels (v1.12.0), ‘Permutation Analysis of Linear Models’ (PALM)^57^ and ‘Network-Based Statistics’ (NBS)^58^ toolboxes, respectively.

Global GM, WM and lobular volumes (% of estimated total intracranial volume (eTIV) to account for differences in head size between participants) were compared between controls and patients using a one-way (GM and WM separately) or multivariate (across GM lobules) ANOVA, as well as a function of NMDAS and mutation load using linear regression analysis. Age and sex effects were accounted for by including them in the model as additional regressors.

For the VBM results and to test for between-group differences, voxel-wise comparisons were performed for GM density and R_1_ maps separately, after which joint inference over the two modalities was performed using Non-Parametric Combination (NPC) and *n*=5000 permutations.^59^ Statistical results were corrected for age, gender and eTIV. Statistical testing was restricted to either GM or WM, as earlier results showed that the m.3243A>G genotype mostly affects GM tissue^16^. Here, the explicit masks were obtained by thresholding (at 0.5) the corresponding SUIT cerebellar probability maps. Finally, after multiple comparison correction (i.e., across voxels and modalities)^60^ using Family-Wise Error (FWE, *q-FWE* = .05) of the statistical T-maps, corresponding clusters of significant differences were exported for visualization and used as additional ROIs for functional connectivity analyses, respectively.

Differences in ROI-to-ROI functional connectivity – defined by the Pearson’s correlation coefficients between a ROI’s across-voxels averaged blood-oxygen-level-dependent (BOLD) timeseries and another ROI’s BOLD timeseries (‘edge’) – between patients and controls, were examined using the NBS statistic, while controlling for age, gender, education and eTIV. Note that the entire connectome (i.e., cortical + cerebellar ROIs) was used at this stage. Cohen’s *d* effect sizes were computed for each significant edge. Multiple regression was used to test for significant correlations of functional connectivity with disease severity and cognitive performance scores across patients only. Bonferroni correction was applied to control for multiple comparisons (i.e., *p* < .05/*n_tests_*).

Finally, summed ROI-based effect size maps (i.e., between groups, as well as those within patients) were decoded into a list of terms to infer mental processes from the observed pattern. To do so, the summed surface-based effect size map was projected back to volume space and smoothed using a gaussian smoothing kernel (*σ* = 2 mm, while ignoring zero-valued voxels) using the ‘metric-to-volume-mapping’, and ‘volume-smoothing’ functions in Connectome Workbench, respectively. A GC-LDA model, in conjunction with results from 14,371 studies within the Neurosynth database, were then used to extract a set of terms. The resulting term’s weight is associated with its relative spatial correspondence with the statistical map’s cortical pattern.^61, 62^

### Data availability

All automatic anatomical and functional data (pre-)processing steps as detailed above have been implemented in a custom and publicly available Snakemake^63^ workflow (https://github.com/royhaast/smk-melas). Raw and processed patient data cannot be made publicly available due to institutional privacy restrictions.

## Results

Example quantitative R_1_ (sec^-1^, left column), cerebellar tissue masks (middle) and density map (a.u., right) for a control subject (top row) and an m.3243A>G patient (bottom row, 24 vs. 38 yrs. old, respectively) are depicted in Fig. 1 across a single sagittal slice. As can be observed, larger inter-folial spaces are visible in the R_1_ (first column) and segmentation images (middle column) for the patient, as indicated by the dashed red lines, compared to the control subject.

**Figure 1.**
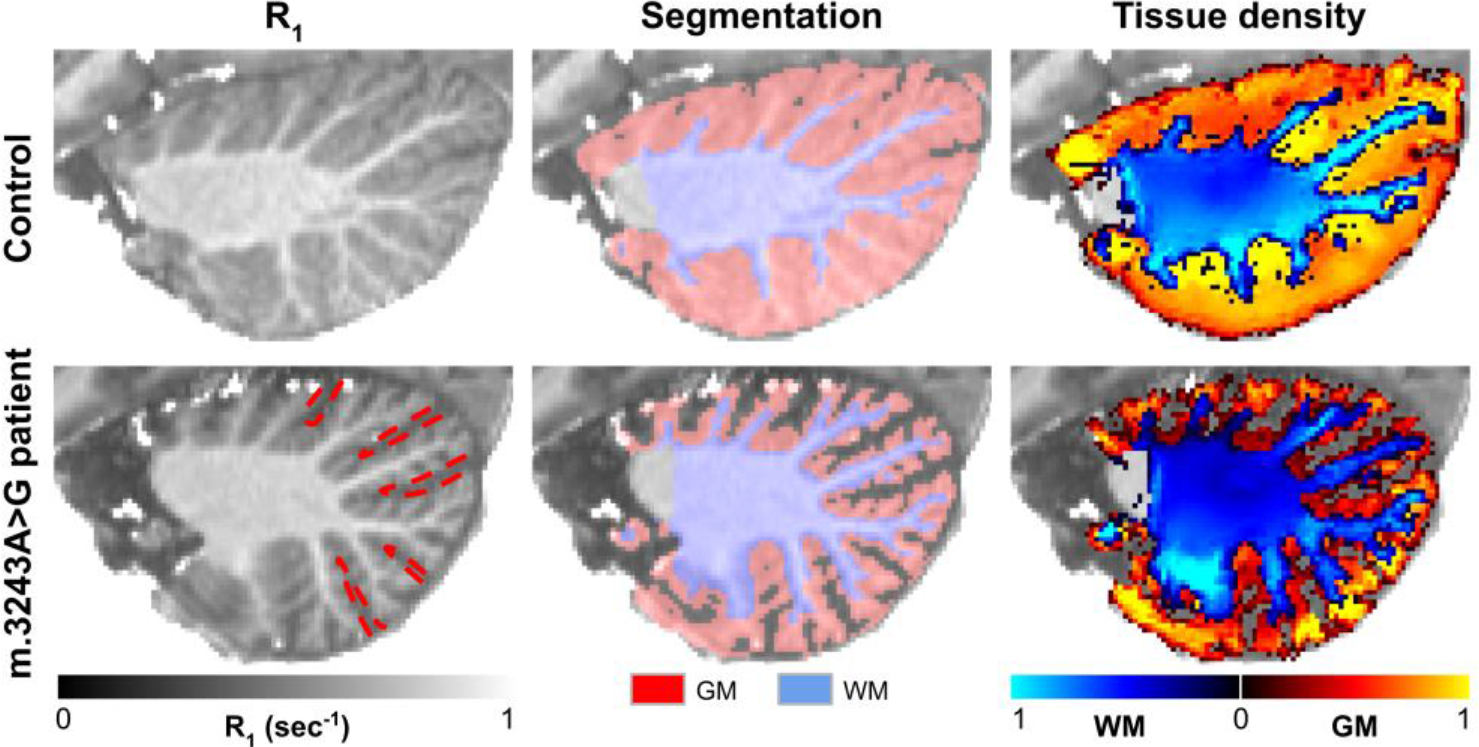
Example data. Left to right: R_1_, GM (red) and WM (blue) segmentation masks, and corresponding tissue density maps are shown for a control (top row) and m.3243A>G patient (bottom row). Dashed red lines indicate the inter-folial spacing for the patient.

Average GM volume was significantly lower for the patient group (*F*_1,68_ = 14.96, *p* < .001, corrected for age, gender and eTIV), while this main effect was negligible for WM (*F*_1,68_ = 0.733, *p* > .05, see red vs. blue dots in top panel in Fig. 2). Significant correlations between average GM, not WM, volume and NDMAS (*p* < .001) and UEC mutation load (corrected for sex, *p* < .001) were observed. This pattern is consistent across (un)corrected heteroplasmy levels in both UEC and blood, while most apparent for the NDMAS section II subscore, and LDST and Stroop cognitive scores (see Supplementary Fig. 1).

**Figure 2.**
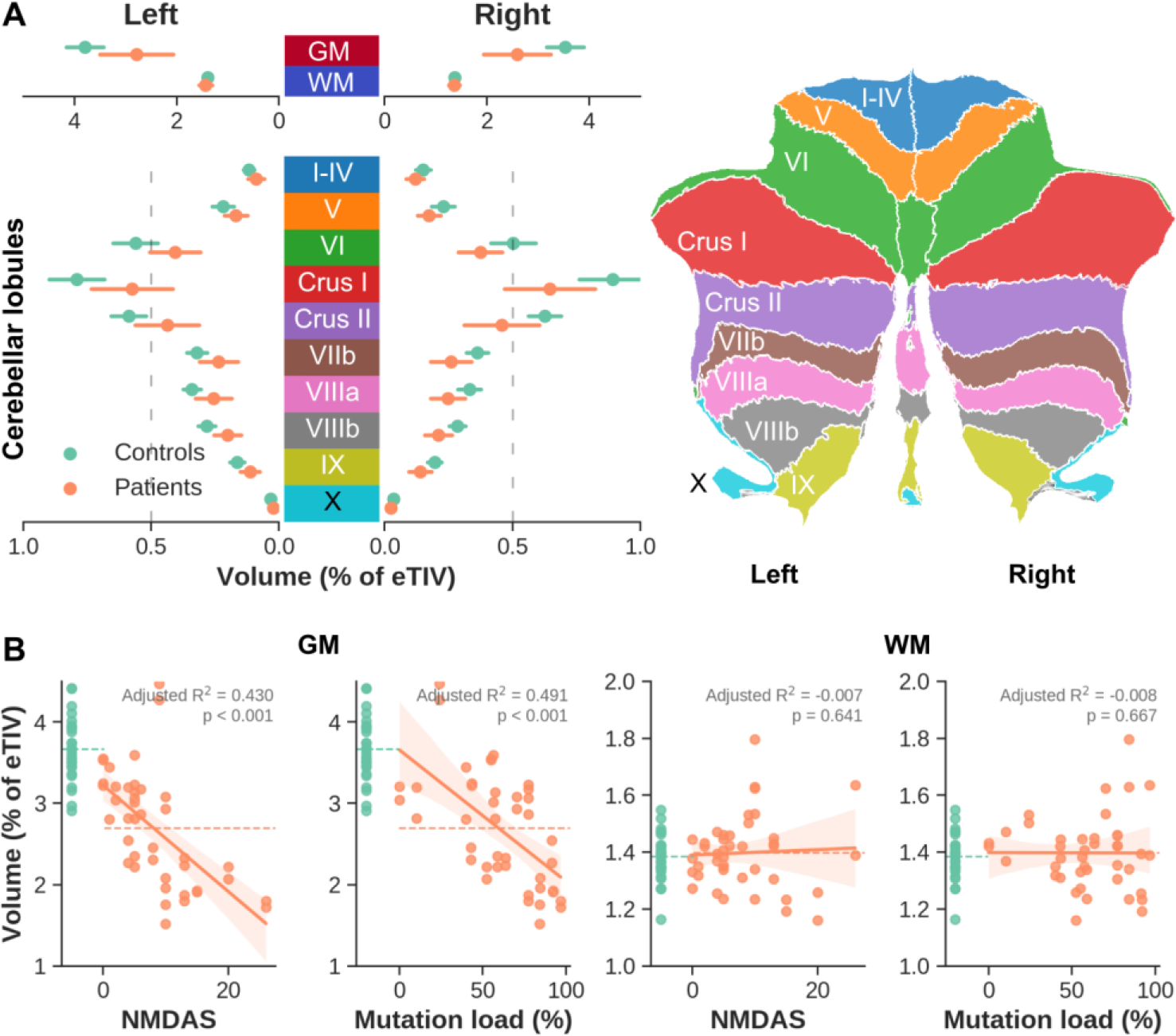
Cerebellar GM and WM volumes. **(A)** Comparison of volume (presented as % of eTIV on the x-axis) between controls (green) and m.3243A>G patients (orange) for left and right hemisphere GM and WM (top), as well as per cerebellar lobule GM (bottom), color-coded based on the right panel legend. **(B)** First two columns: correlation between GM volume (y-axis) and NMDAS or corrected UEC mutation load (x-axes). Last two columns: similar to first two columns but using WM volume (y-axis).

More detailed, voxel-wise comparison of GM density and R_1_ in Fig. 3 were used to better describe the spatial-specificity of volumetric differences between groups. Both modalities were tested individually and then combined for joint inference using Fisher’s NPC to extract significant clusters. Note that results are visualized on a flat representation of the cerebellum but that the analyses were performed in volume space (see Supplementary Fig. 2 for the volume to flat representation correspondence). GM density was found consistently higher for control subjects (i.e., in red), while differences in R_1_ are more variable but revealing a similar pattern with higher R_1_ for the controls. A total of eight clusters of voxels characterized by significant differences in both GM density and R_1_ (Fisher combined *p_permuted_* < .05, delineated by solid back lines) were extracted. The six largest clusters (1-6), characterized by a symmetric distribution across left and right hemispheres (see also 3D rendering), were selected for further characterization using public atlases as well as *in vivo* resting-state fMRI data.

**Figure 3.**
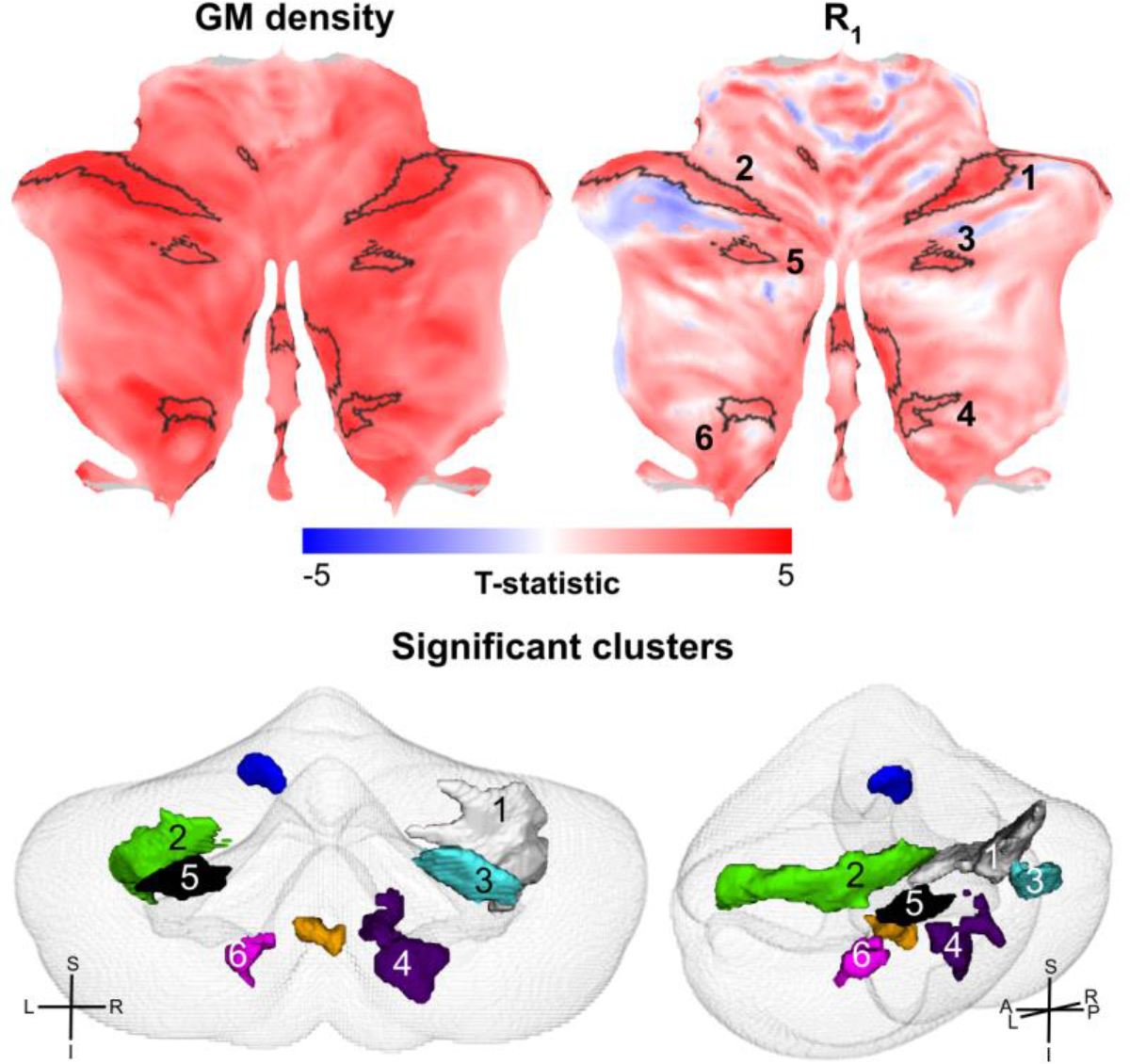
Voxel-based statistical results. Flatmap representation of the statistical result when comparing GM density (left) and R_1_ (middle) maps between controls and patients. Significant clusters after joint interference are delineated using solid black lines on the flatmaps and represented as 3D meshes (bottom). Orientation crosses provide references to left-right (L-R), superior-inferior (S-I) and anterior-posterior (A-P) axes.

First, to evaluate whether the significant clusters tend to colocalize with predefined anatomical (or functional) parcels, we quantified cluster sizes and their overlap for each cluster–parcel combination (e.g., cluster 1 vs lobule VI, Fig. 4, left panel). Here, the dashed black line represents the individual cluster sizes (in number of voxels, sorted from largest to smallest) while the stacked bar plot indicates the proportion (%) of each cluster that falls within the respective color-coded atlas region (see middle panel). The two largest clusters (i.e., one and two, covering 1,960 and 1,266 mm^3^, respectively) were equally positioned across lobule VI (48.31 and 40.36 % of their total volume, respectively) and Crus I (51.96 and 59.64 %). Cluster sizes drop strongly from cluster 3 with volumes decreasing from 427 to 166 mm^3^. Taken together (right panel), lobule VI (32.01 %) and Crus I (50.97 %) show the largest overlap with all clusters.

**Figure 4.**
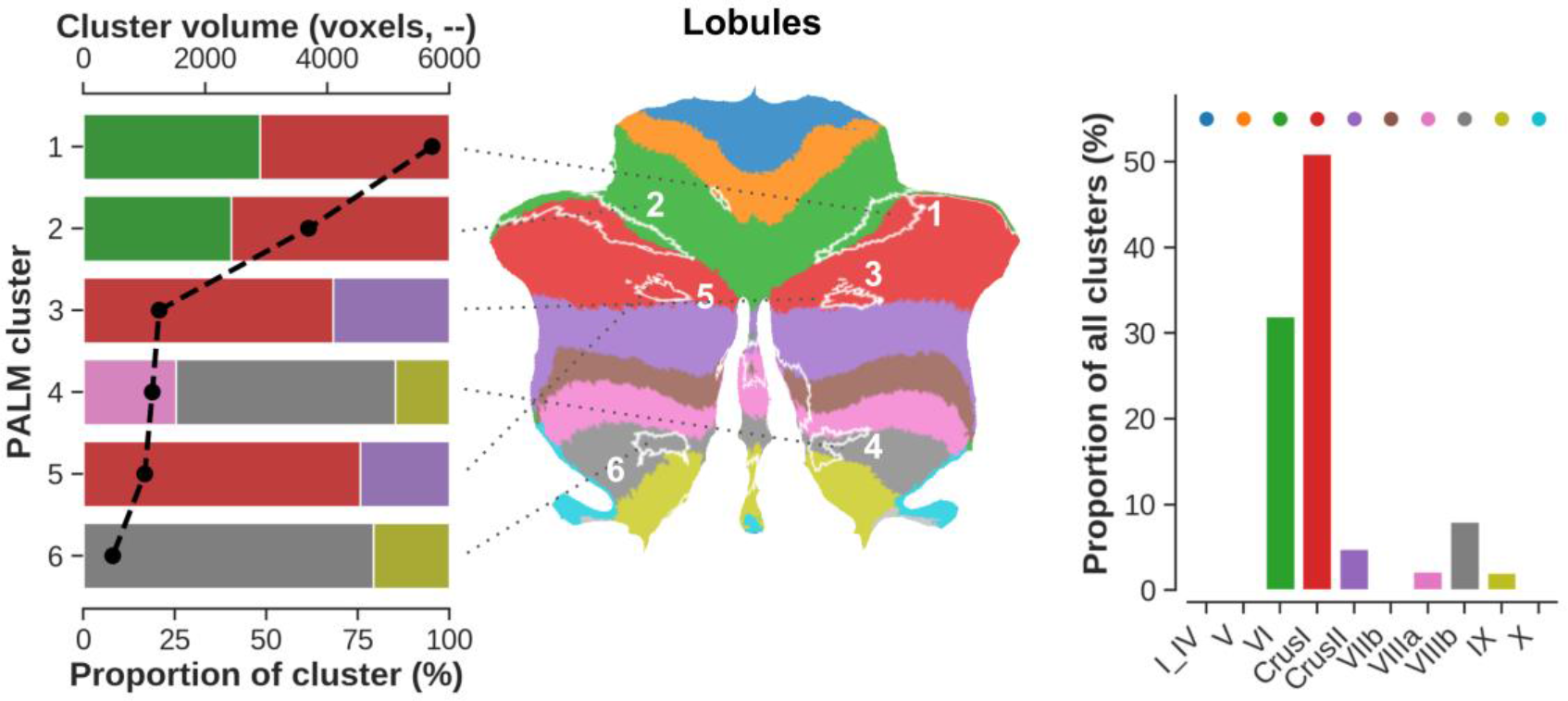
Spatial distribution of the significant clusters with respect to the cerebellar lobules. Left to right: stacked bar plot showing statistical (i.e., PALM) clusters (y-axis), ordered from largest at the top (cluster 1) to smallest at the bottom (cluster six, in voxels, top x-axis). Here, the width of each individually-colored bar represents the proportional overlap (bottom x-axis) with the respective lobule. For example, 50 % of cluster one overlaps with Crus I. Middle panel shows a flatmap representation to visualize the localization of each cluster across the cerebellar GM with respect to its lobules. Right panel shows the proportional overlap (y-axis) across all clusters per lobule (x-axis). For example, 50 % of significant voxels fall within Crus I.

Functionally (see Supplementary Fig. 3), clusters one (74.36 %) and two (72.48 %) strongly colocalize with FPN. Overall, most voxels characterized by a significant difference in GM density and R_1_ between groups lie within FPN (52.57 % of total voxels), followed by the DMN (26.21 %), VAN (13.96 %) and SMN (6.66 %), while the overlap with visual, DAN and limbic networks remain negligible (i.e., < 1%).

Second, to characterize the functional signatures of the affected tissue, connectivity profiles (i.e., ROI–ROI functional timeseries correlation) extracted from *in vivo* rs-fMRI data were explored and compared between groups. Example rs-fMRI cortical and cerebellar data for a control subject and m.3243A>G patient for a corresponding brain coactivation timepoint (i.e., DMN) are shown in Fig. 5A. Subsequent statistical comparison between groups revealed one network across the cortical and cerebellar nodes with 167 edges that were characterized by a significant reduction in connectivity strength for the m.3243A>G patients. Supplementary Fig. 4 shows the statistical and corresponding significance matrices. Across all 167 edges, 63 edges (37.72 %, solid black lines in Fig. 4B) showed a significantly impacted (*p* < .05, NBS corrected, controls > patients) connectivity strength between the cerebellar clusters and a cortical ROI. No difference was observed between left and right cerebral hemispheres in their connectivity to the cerebellum in patients (*F*_1,49_ = 0.008, *p* > .05). See Supplementary Figs. 5 and 6 for the cluster-wise Cohen’s *d* effect sizes and cortical connectivity profiles, respectively. Taken together, these affected cortical ROIs (delineated using a solid black line in Figs. 5C and D) are predominantly positioned along a lateral parietal to frontal band where most prominent group effects are observed in the (especially left hemispheric) frontal regions with cortical ROI’s characterized by reduced connectivity with at least two cerebellar clusters (Fig. 5C). In parallel, the m.3243A>G mutation most significantly impacts the frontal regions (Fig. 5D).

**Figure 5.**
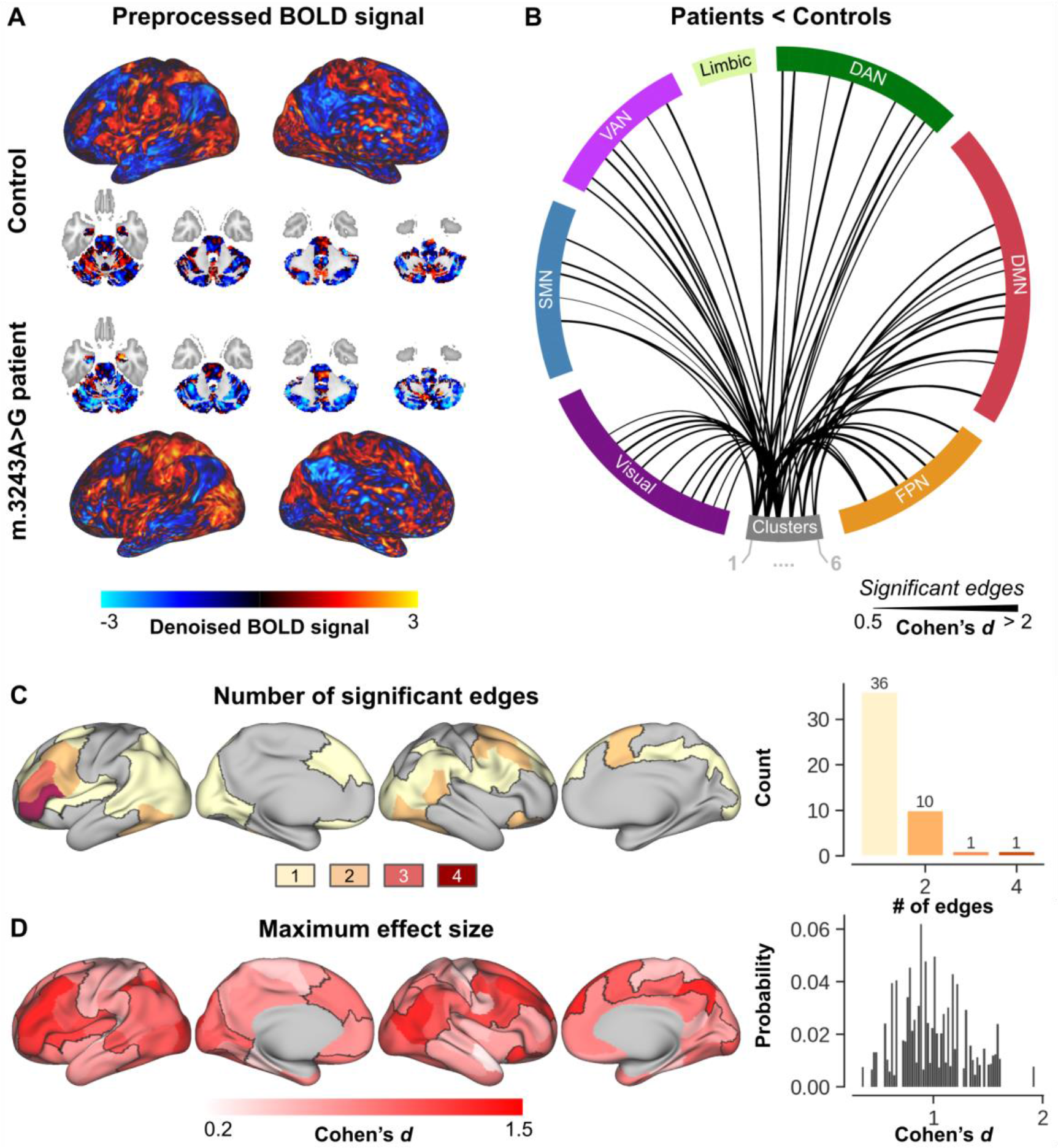
Characterization of cerebello–cortical functional connectivity. **(A)** Visual comparison of the denoised rs-fMRI cortical and cerebellar data for a control subject (top part) and m.3243A>G patient (bottom part) at a corresponding brain coactivation timepoint. **(B)** Significantly reduced (solid black lines) cerebello–cortical (separated per large-scale brain network) connections in m.3243A>G patients compared to controls. **(C)** Surface-wise visualization of the total number of significantly reduced edges (in m.3243A>G patients) per cortical ROI. For example, a cortical ROI will be colored yellow if it shows reduced connectivity to only a single cerebellar cluster, but red if it shows reduced connectivity to four out of the six clusters. ROIs not affected at all are shown in gray. **(D)** Corresponding maximum effect size per ROI. Briefly, all ROIs are characterized by six T-statistical values, based on the group-wise difference for each of the cerebellar clusters. The maximum is then mapped onto the cortical surface.

**Figure 6.**
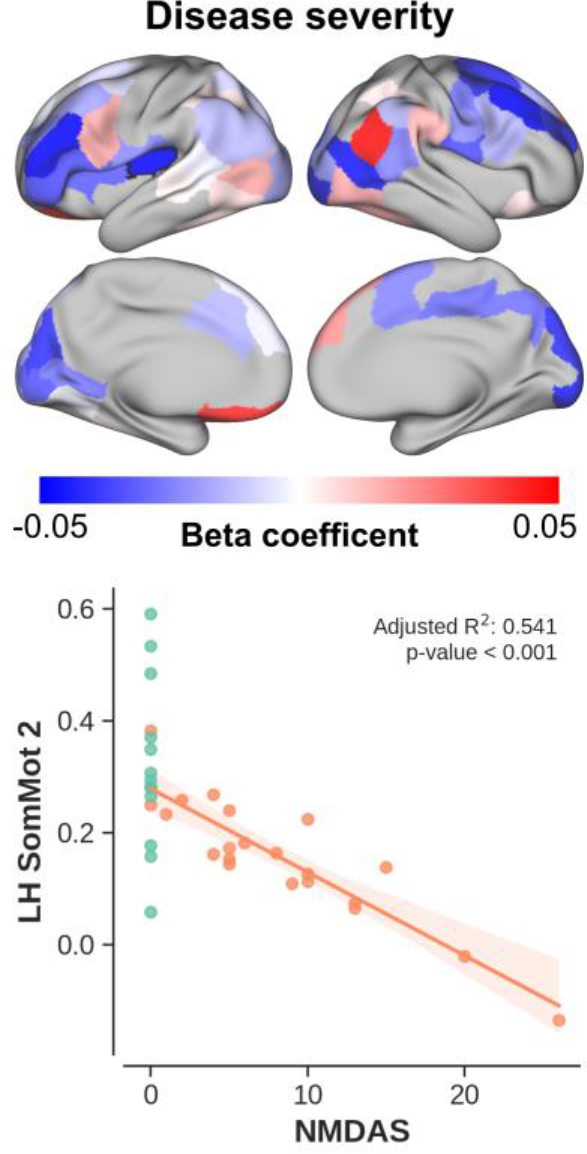
Disease severity vs connectivity. Top panel: beta coefficients (i.e., explained change in connectivity strength per unit change in NMDAS) per cortical region mapped onto the cortical surface. Bottom panel: scatter plot showing the change in connectivity (for patients, in orange) as function of NMDAS for the cerebello–cortical pair characterized by the strongest correlation. Control data are shown for comparison (in green).

Once we identified the edges that were statistically reduced in the patient group, correlation analyses were used to investigate whether the observed effect was stronger in patients characterized by (1) a more severe disease phenotype (Fig. 6) or (2) worse cognitive performances (Fig. 7). Overall, but not exclusively, functional connectivity scales negatively with increasing NMDAS score (i.e., more severe phenotype) across patients. Again, this effect is strongest at the frontal lobe, as well as the insular cortex. For example, a negative correlation (*p* < .001) is visible between NMDAS and cerebellar functional connectivity to a region embedded within the SMN (outlined with a black solid line in upper left surface-based display). Positive correlations are observed across several regions too. However, in contrast to the negative correlations, these are spread asymmetrically across the brain, without a strong spatial preference.

**Figure 7.**
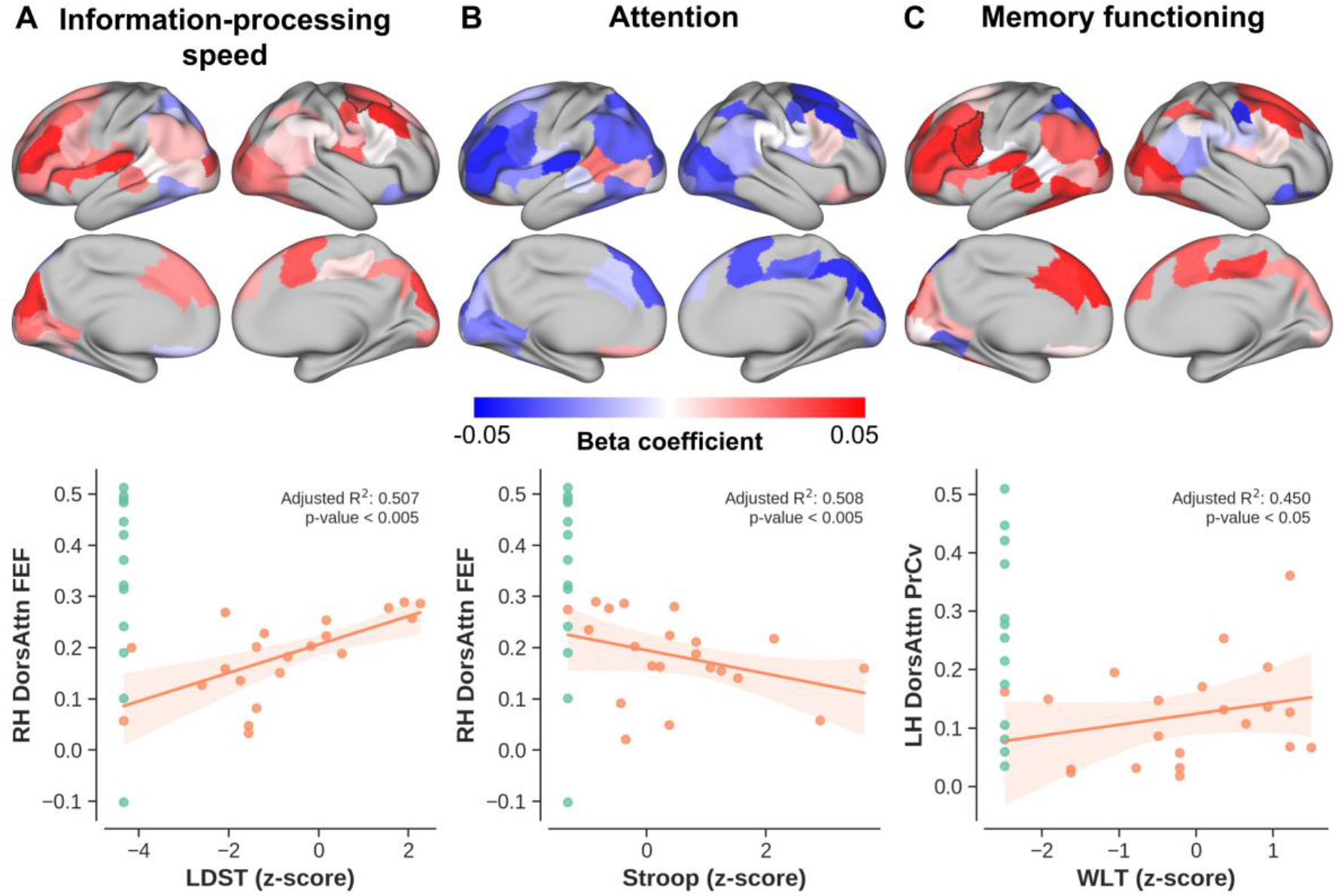
Cognition vs connectivity. Similarly to Figure 6 with top panels showing the beta coefficients (i.e., explained change in connectivity strength per unit change in cognitive test score) per cortical region and bottom panels showing scatter plot with the change in connectivity as function of **(A)** LDST, **(B)** Stroop and **(C)** WLT corresponding to information-processing speed, attention and memory functioning, respectively.

Functional connectivity decreases with decrease in cognitive performance based on the patients’ letter-digit substitution task (LDST, i.e., higher is better, see Fig. 7A for the corresponding cortical ROI beta coefficients), Stroop (i.e., higher is worse, Fig. 7B) and 15 words-learning task (WLT, higher is better, Fig. 7C) test scores. This effect is most consistent across regions for LDST (i.e., information processing speed) and Stroop (attention), but more variable for WLT (memory).

Group-wise, disease severity and cognitive performance effect sizes (see Supplementary Fig. 7A for their comparison) were summed to identify cortical regions characterized by the most consistent change in their functional connectivity with the cerebellar clusters. Summed effect sizes ranged from 4.31 in parietal regions up to 16.99 in frontal regions (Supplementary Fig. 7B). Comparison of the corresponding spatial pattern to the results extracted from 14,371 studies in the Neurosynth database revealed a strong correlation with broad terms such as ‘visual’ (‘correlation weight’ = 10097.54, Supplementary Fig. 7C), ‘motor’ (5161.05) and ‘attention’ (3906.84) where the term’s font size scales with its corresponding weight.

## Discussion

The m.3243A>G genotype is characterized by a large phenotypic spectrum across patients.^2, 4^ In this work, we employed the most detailed MRI dataset available in a relatively large population of patients carrying the m.3243A>G mutation to define alterations of the spatial pattern of cerebellar macro- and microstructural features, as well as their functional connectivity to cortical areas.

### Impact on cerebellar structure

In line with our earlier cerebral cortical findings,^16^ the current results show that the m.3243A>G mutation induced (almost exclusively) cerebellar GM tissue changes. Cerebellar GM atrophy worsened with increased severity based on the NMDAS score as well as a higher mutation load measured in both blood and urine epithelial cells similar to that observed for the cerebral cortex. This follows previous *in vivo* and *ex vivo* observations by means of a higher degree of abnormal radiotracer binding^15^ and neuronal loss^17^ in cerebellar tissue from more severely affected patients, respectively. The GM density changes were accompanied by a decrease in R_1_, indicating a reduced concentration of intracortical myelin and iron.^33^ In contrast, the WM tissue remained unaffected, independent of disease severity based on both clinical phenotype and mutation load. Together these suggest that the GM tissue’s integrity can become severely impaired in m.3243A>G patients, compared to a group of controls. While this effect appeared to be global (i.e., across the entire GM), statistical testing revealed several ‘hot spots’, or clusters, spread across the cerebellar lobules in a systematic left vs. right fashion for the largest clusters. Spatial characterization of these clusters with respect to a cerebellar anatomical atlas and its lobulation^45^ revealed a strong bias towards lobules VI and Crus I, harboring almost 80% of all the significant voxels. In the following, we will contextualize these results using the relevant literature, focusing mostly on the interplay between mitochondrial (dys)functioning and neuronal integrity.

As mentioned in the Introduction, the cerebellum is known for its immensely folded structure that accounts for the majority of the neuronal cell bodies found in the brain. It covers a total area of about 1,590 cm^2^ when unfolded, rendering it considerably more dense compared to the roughly 2,000 cm^2^ area of the eight times volume of the larger cerebral cortex.^19, 64^ Consequently, the cerebellar tissue requires a steady and relatively vast supply of nutrients (mostly carbohydrates and fatty acids) to nourish the basal level of activity of its densely packed neurons.^65^ The metabolic processes to release the stored energy from these nutrients and generate ATP, the actual energy substrate, is coregulated by a collection of respiratory chain subunits located within the mitochondria.^66^ As such, mitochondrial mutations, like the one central to this work, will lower the mitochondria’s efficiency to produce ATP through oxidative phosphorylation^67^ and affect the functioning of multiple organs, when crossing a tissue-specific threshold.^68^ Below the threshold, the mutation remains unnoticed. It has been shown in myoblasts (i.e., embryonic progenitor cells that give rise to muscle cells) from a single MELAS patient that having a >80-90% m.3243A>G mutation load leads to impaired translation of all mitochondrial encoded respiratory chain subunits with a decrease in ATP synthesis as result.^69^ Recent work has confirmed this observation in human neurons using induced pluripotent stem cell (iPSC) technology.^70^ Additionally, the authors observed differences between low and high levels of heteroplasmy iPSC neurons’ anatomy where high levels (71%) of the m.3243A>G mutation appeared to reduce synapses, mitochondria, and dendritic complexity. This is in line with earlier work that linked mitochondrial dysfunction, as well as reduced mitochondrial mass, with altered neuronal dendritic morphology and remodeling in vitro and *in vivo*, including direct measurements in the cerebellum.^17, 70, 71^ Additionally, simulations based on a m.3243A>G biophysical model suggest that cell volume decreases with increasing heteroplasmy to prevent potential energy crises^72^ while the absolute number of mitochondria is often increased in m.3243A>G patients.

Moreover, biochemical deficits and clinical implications only appear once the patient’s heteroplasmy level surpasses a certain cellular or tissue-specific threshold.^1, 67, 73^ As we only included patients, this implies that the threshold at least in some tissues were surpassed, although symptoms could be very subtle and with cerebellar GM volumes similar to those in the lower regime of observations across healthy controls. The linearly (and significantly) decreasing GM volume as a function of mutation load is indicative of an additional gradual effect of the genotype on the cerebellar tissue changes once the threshold of expression is surpassed (i.e., more profound enzyme deficiency). It is important to note that a similar, linear relationship was observed when opposing the volumetric measures to the NMDAS score. Patients with a more severe disease phenotype appear to be characterized by the strongest atrophy. Nevertheless, the heterogeneity and complexity of the m.3243A>G phenotype challenges theoretical understanding of their causation and requires longitudinal tracking of disease progression.

Taken together, the observations discussed above strongly suggest that the m.3243A>G mutation specifically impacts the GM tissue through neuronal morphological changes. Here, our spatial characterization using voxel-wise analyses – that showed a bias towards lobules VI (∼30 %) and Crus I (∼50 %), located along the superior-posterior portion of the cerebellum – might be used to further deduce the anatomical specificity of these changes towards specific cytoarchitectonic, molecular and/or structural connectivity features.^74^

Cytoarchitectonically, the cerebellar GM is characterized by a distinct (i.e., compared to the neocortex), uniform three-layer architecture composed of the inner granular, outer molecular layer and in between a sheet of Purkinje cells which are solely responsible for directing information away from the cerebellum.^75^ Independent of lobulation, ‘transversal zones’ have been identified by leveraging the molecular topography defined by the expression of specific genes across the cerebellum. Interestingly, most significant voxels lie within a central zone characterized by Purkinje cells expressing zebrin II,^76^ which is analogous to aldolase C72,^77^ an important player in glycolytic ATP biosynthesis,^78^ posing an indirect link to mitochondrial dynamics.^79^ While spinocerebellar ataxia seems to involve neurodegeneration of motor-related cerebellar regions,^80^ m.3243A>G-related atrophy might be restricted to certain Purkinje subtypes (e.g., zebrin II+). However, the molecular characterization remains a complex issue and out of the scope of this manuscript. In parallel, the cerebellar cortex can be parcellated based on its anatomical connectivity. In contrast to the transversal zones based on genetic markers, these zones run in a longitudinal fashion, perpendicular to the long axis of the lobules. Most significant voxels lie within zones that appear to receive input from the principal olive nucleus. However, the current results do not show a clear bias towards a specific (set of) zone(s) with the significant clusters spanning from the lateral hemispheres up to the (para)vermis. More coarsely, tracer studies in the macaque monkey show a distinction between prefrontal (mainly lobules Crus I and II) and motor (all other) modules, with anatomical connections running to the respective cortical areas.^23, 81^ With Crus I being the most affected lobule, especially prefrontal connectivity might be impacted.^82^ However, *in vivo* fMRI data is necessary to characterize the functional consequences, which will be discussed next.

### Impact on cerebellar functional connectivity

It is the growing consensus, supported by electrophysiological mapping in a range of species, that the cerebellum’s functional modules are not shaped by its lobules but extend beyond its fissures.^74^ Drawing conclusions solely based on comparisons with previously published anatomical parcellations and literature might therefore paint an incomplete picture. As such, we leveraged an openly available functional parcellation, as well as acquired rs-fMRI data to more precisely map out the impact of the observed differences on the brain’s functioning, and potential correlations with the clinical phenotype, based on disease severity and cognitive performance.

Several studies have used the synchronization of rs-fMRI signals between brain regions to identify seven large-scale brain networks.^56, 83^ From a historic perspective, the function of the cerebellum has been linked to the sensorimotor system. However, the cerebellum appears to play an important role across multiple of the identified large-scale cortical brain networks.^84, 85^ Our results show great overlap with cerebellar fractions of four of these identified networks but most prominently with FPN (>50 %), followed by DMN (∼25 %) and VAN (∼15 %). All regions that show functional connectivity with associative regions of the cerebral cortex (and found to be similarly affected in schizophrenic patients).^86^ The FPN, also known as the ‘central executive network’, plays an important role in higher cognitive functions by actively maintaining and manipulating information in working memory, for rule-based problem solving and for decision making in the context of goal-directed behavior.^87^ Unlike all other networks, the FPN is disproportionately (i.e., ∼two-fold) expanded in the cerebellum compared to the cerebral cortex and might therefore play a relatively important role at the whole-brain scale.^84, 88^ Damage to the FPN in the cerebellum disturbs a broad range of control functions, including task switching, working memory retrieval, visuo-spatial integration, language, and an overall reduction in intellectual function,^89^ collectively known as the cerebellar cognitive affective syndrome.^90^ Cognitive deficits are not uncommon in mitochondrial disorders and prevalent in up to a third of m.3243A>G patients.^5, 68^ While cognitive performance appears to reduce in general, distinct domains, including verbal comprehension, perceptual reasoning, working memory, processing speed, and memory retrieval, were found to be affected in particular.^91^ Similarly, the lower LDST and Stroop test scores indicate impaired information processing speed and attention in the current cohort of patients. In both cases, adequate performance thrives on the fluent selection of relevant visual features through neuronal computations in frontal, parietal, and/or limbic areas that are then projected to occipital (i.e., visual) areas.^92, 93^

Additionally, we used rs-fMRI data to identify impaired brain networks in our patients. Prior evidence is scarce and only one study has systematically investigated changes in the whole brain’s functional topology of m.3243A>G patients.^94^ Here, modularity analysis (e.g., network efficiency) revealed that patients had altered intra- or inter-modular connections in default mode, frontoparietal, sensorimotor, visual and cerebellum networks. Our results – using analyses that were particularly focused on the interplay between the affected cerebellar clusters and the rest of the brain – revealed a single network of regions that showed significantly reduced connectivity in the m.3243A>G patients. Spatial characterization of this network shows a strong emphasis on frontal and parietal lobe regions with especially the (left) frontal lobe characterized by impaired connectivity with the cerebellum (e.g., based on the number of significant edges) that intensifies in the more severely affected patients, based on the NMDAS score. This bias towards the frontal lobe, also known as fronto-cerebellar dissociation, has been found to increase the difficulty for a person to select the appropriate response to a stimuli, or to initiate the response (i.e., executive functioning).^95^ Moreover, focal frontal and parietal lobe lesions resulted in increased errors and slowness in response speed during the Stroop test.^96, 97^ Similarly, the frontal-parietal cortical network appears to be strongly engaged during the LDST task.^98^ In line with these previous studies, our correlational analyses between functional and cognitive profiles show that cerebello–cortical connections characterized by a significant group effect, are weaker in patients with lower LDST and Stroop performances. Additionally, the left frontal lobe is considered the anterior convergence zone of the dorsal (i.e., phonology) and ventral (i.e., semantics) language streams,^99^ thus playing an essential role in this dual-stream model. The central role of the frontal lobe in this model of language processing explains the appearance of terms like ‘language’, ‘words’ and ‘semantic’ when comparing our statistical maps to those included in the NeuroSynth database^62^ and could provide novel insights into the cognitive deficits related to the m.3243A>G mutation, and/or mitochondrial diseases in general.

### Clinical implications

The clinical manifestation of the m.3243A>G mutation is characterized by a wide variability in nature and severity of symptoms.^4^ In a small subset of carriers the mutation induces a severe phenotype, such as the MELAS syndrome, with stroke-like episodes, encephalopathy and progressive cognitive difficulties.^4, 7^ Current results – based on mildly affected patients with relatively low Barthel and NDMAS scores – are therefore most relevant for more common manifestations (e.g., MIDD and myopathy), and patients characterized by a mutation load range like the current study population, while generalizability to more severe m.3243A>G clinical phenotypes is lower. Regardless, cerebellar integrity, in particular the subregions identified by the current work, could serve as a target for longitudinal disease tracking (e.g., to study brain-phenotype relationship) and/or evaluate the efficacy of potential treatments (e.g., L-Arginine supplementation) across the entire spectrum of patients.^100^ Based on current and previous^16^ findings, structural changes in m.3243A>G patients range from large-scale deformations (e.g., enlarged ventricles) to fine-scale (e.g., local tissue T1) changes, depending on the severity of the case. While ventricular volume changes can readily be detected at conventional field strengths (i.e., ≤ 3T), the use of 7T MRI might be crucial to detect the subtle differences in structures like the cerebellum. The steady increase in the number of clinically approved 7T MRI scanners will increase feasibility to apply these methodologies in more clinical-oriented applications (e.g., diagnosis, drug development).

### Technical considerations

Despite the advantage of using high-resolution anatomical and functional data, the cerebellum’s fine-scale anatomy might introduce signal contamination.^19^ Partial voluming effects (in regions characterized by a thin cortex) between the GM, WM and CSF voxels’ fMRI timeseries, in particular, will affect downstream functional connectivity analyses. We counteracted this at four different stages. First, during tissue segmentation by careful isolation of the cerebellar tissue. Second, during fMRI data preprocessing, by using a one-step resampling (and thus interpolation) procedure. See also Supplementary Fig. 8 for the residual but negligible impact of this step on the volume, R_1_ and functional connectivity results. Third, during fMRI signal denoising, by regressing out WM and CSF signal timeseries at the voxel-level. Finally, by modeling whole-brain functional connectivity as a graph during statistical analyses using the NBS, based on data from the entire study population.^58^ Together, these rendered the identified significant network minimally sensitive to cerebellar ROI- and/or patient-specific outliers.

### Conclusions

In summary, the current results indicate that the m.3243A>G mutation significantly impacts the cerebellum with strongest changes observed in most severely affected patients, based on genetic, clinical and cognitive features. The impact of the m.3243A>G mutation ranges from reduced GM tissue integrity to impaired functional connectivity with cortical brain regions. Spatial characterization reveals that these changes occur especially in tissue and regions related to the FPN, crucial for information processing speed and selective attention. Combined with our previous work,^16^ it provides insight into the neuropathological changes and a solid base to guide longitudinal studies aimed to track disease progression.

## Acknowledgements

We would like to thank the patients and control participants who agreed to take part in this study. In addition, we thank Rutger JT IJsselstein, Suzanne CEH Sallevelt, Florence van Tienen and Elia Formisano for their technical assistance and/or constructive feedback.

