## Supplementary material for "Neurodegenerative and functional signatures of the cerebellar cortex in m.3243A>G patients"

**Supplementary Table 1 Individual NMDAS scores.** Patient-specific compound NDMAS scores for total, subsections I ('current function'), II ('system-specific involvement') and III ('current clinical assessment'), as well as individual scores for each of the neurological features. Median (IQR) values are provided in the last column.

| | **1** | **2** | **3** | **4** | **5** | **6** | **7** | **8** | **9** | **10** | **11** | **12** | **13** | **14** | **15** | **16** | **17** | **18** | **19** | **20** | **21** | **22** | **Median (IQR)** |
|---|---|---|---|---|---|---|---|---|---|---|---|---|---|---|---|---|---|---|---|---|---|---|---|
| **Overall** | | | | | | | | | | | | | | | | | | | | | | | |
| Total | 20 | 1 | 5 | 10 | 10 | 17 | 5 | 5 | 13 | 26 | 4 | 6 | 8 | 10 | 4 | 0 | 9 | 0 | 2 | 5 | 13 | 15 | 7 (8) |
| Section I – Current function | 9 | 0 | 0 | 2 | 1 | 11 | 0 | 0 | 6 | 6 | 1 | 1 | 1 | 4 | 3 | 0 | 4 | 0 | 0 | 3 | 2 | 5 | 1.5 (4) |
| Section II – System-specific involvement | 6 | 0 | 4 | 8 | 8 | 5 | 5 | 5 | 6 | 14 | 2 | 5 | 3 | 6 | 1 | 0 | 1 | 0 | 2 | 1 | 11 | 5 | 5 (4.75) |
| Section III – Current clinical assessment | 5 | 1 | 1 | 0 | 1 | 1 | 0 | 0 | 1 | 6 | 1 | 0 | 4 | 0 | 0 | 0 | 4 | 0 | 0 | 1 | 0 | 5 | 1 (1) |
| **Neurological** | | | | | | | | | | | | | | | | | | | | | | | |
| Total | 4 | 1 | 1 | 1 | 1 | 3 | 5 | 5 | 3 | 9 | 2 | 1 | 2 | 3 | 3 | 0 | 2 | 0 | 2 | 3 | 5 | 3 | 2.5 (2) |
| Hearing loss | 2 | 0 | 0 | 1 | 1 | 2 | 0 | 0 | 2 | 2 | 1 | 1 | 1 | 3 | 2 | 0 | 1 | 0 | 0 | 2 | 2 | 1 | 1 (2) |
| Psychiatric | 0 | 0 | 0 | 0 | 0 | 0 | 0 | 0 | 0 | 2 | 0 | 0 | 0 | 0 | 0 | 0 | 0 | 0 | 0 | 0 | 0 | 0 | 0 (0) |
| Migraine | 0 | 0 | 0 | 0 | 0 | 0 | 5 | 5 | 1 | 1 | 1 | 0 | 1 | 0 | 1 | 0 | 1 | 0 | 1 | 1 | 3 | 0 | 0.5 (1) |
| Seizures | 0 | 0 | 1 | 0 | 0 | 1 | 0 | 0 | 0 | 0 | 0 | 0 | 0 | 0 | 0 | 0 | 0 | 0 | 1 | 0 | 0 | 0 | 0 (0) |
| Encephalopathic episodes | 0 | 0 | 0 | 0 | 0 | 0 | 0 | 0 | 0 | 0 | 0 | 0 | 0 | 0 | 0 | 0 | 0 | 0 | 0 | 0 | 0 | 0 | 0 (0) |
| Stroke-like episodes | 0 | 0 | 0 | 0 | 0 | 0 | 0 | 0 | 0 | 0 | 0 | 0 | 0 | 0 | 0 | 0 | 0 | 0 | 0 | 0 | 0 | 0 | 0 (0) |
| Cerebellar ataxia | 2 | 1 | 0 | 0 | 0 | 0 | 0 | 0 | 0 | 2 | 0 | 0 | 0 | 0 | 0 | 0 | 0 | 0 | 0 | 0 | 0 | 2 | 0 (0) |
| Neuropathy | 0 | 0 | 0 | 0 | 0 | 0 | 0 | 0 | 0 | 2 | 0 | 0 | 0 | 0 | 0 | 0 | 0 | 0 | 0 | 0 | 0 | 0 | 0 (0) |
| Pyramidal involvement | 0 | 0 | 0 | 0 | 0 | 0 | 0 | 0 | 0 | 0 | 0 | 0 | 0 | 0 | 0 | 0 | 0 | 0 | 0 | 0 | 0 | 0 | 0 (0) |
| Extrapyramidal | 1 | 0 | 0 | 0 | 0 | 1 | 0 | 0 | 0 | 0 | 0 | 0 | 0 | 0 | 0 | 0 | 0 | 0 | 0 | 0 | 0 | 0 | 0 (0) |

**Supplementary Table 2 Data acquisition parameters**

| | $T_1$ | | **Rs-fMRI**[a] |
|---|---|---|---|
| | *3D MP2RAGE* | *3D Sa2RAGE* | *2D MB-EPI* |
| TR, ms | 5000 | 2400 | 2000 |
| TE, ms | 2.47 | 0.78 | 18.8 |
| $TI_1/TI_2$ ($TD_1/TD_2$), ms | 900/2750 | 58/1800 | - |
| Flip angle(s) | 5/3 | 4/10 | 80 |
| Partial Fourier | 6/8 | 6/8 | 6/8 |
| Phase-encoding | A-P | A-P | A-P |
| GRAPPA[b] | 3 | 2 | 3 |
| Reference lines | 24 | 24 | 54 |
| Number of slices (/vols) | 240 sagittal | 88 sagittal | 80 (/300) |
| Field of view, mm | 224 × 224 | 256 × 256 | 198 × 198 |
| Matrix size, mm | 320 × 320 × 240 | 128 × 128 × 96 | 142 × 142 × 112 |
| Acquisition time, m:s | 8:02 | 2:16 | 10:20 |

[a]Five reverse phase-encoded rs-fMRI volumes were acquired to correct for readout-related geometrical distortion. Except for the phase encoding direction (P-A) and number of vols (5), all acquisition parameters were identical. [b]GRAPPA was applied in the phase-encoding direction. Abbreviations: TR = repetition time; TE = echo time; TI = inversion time; TD = delay time.

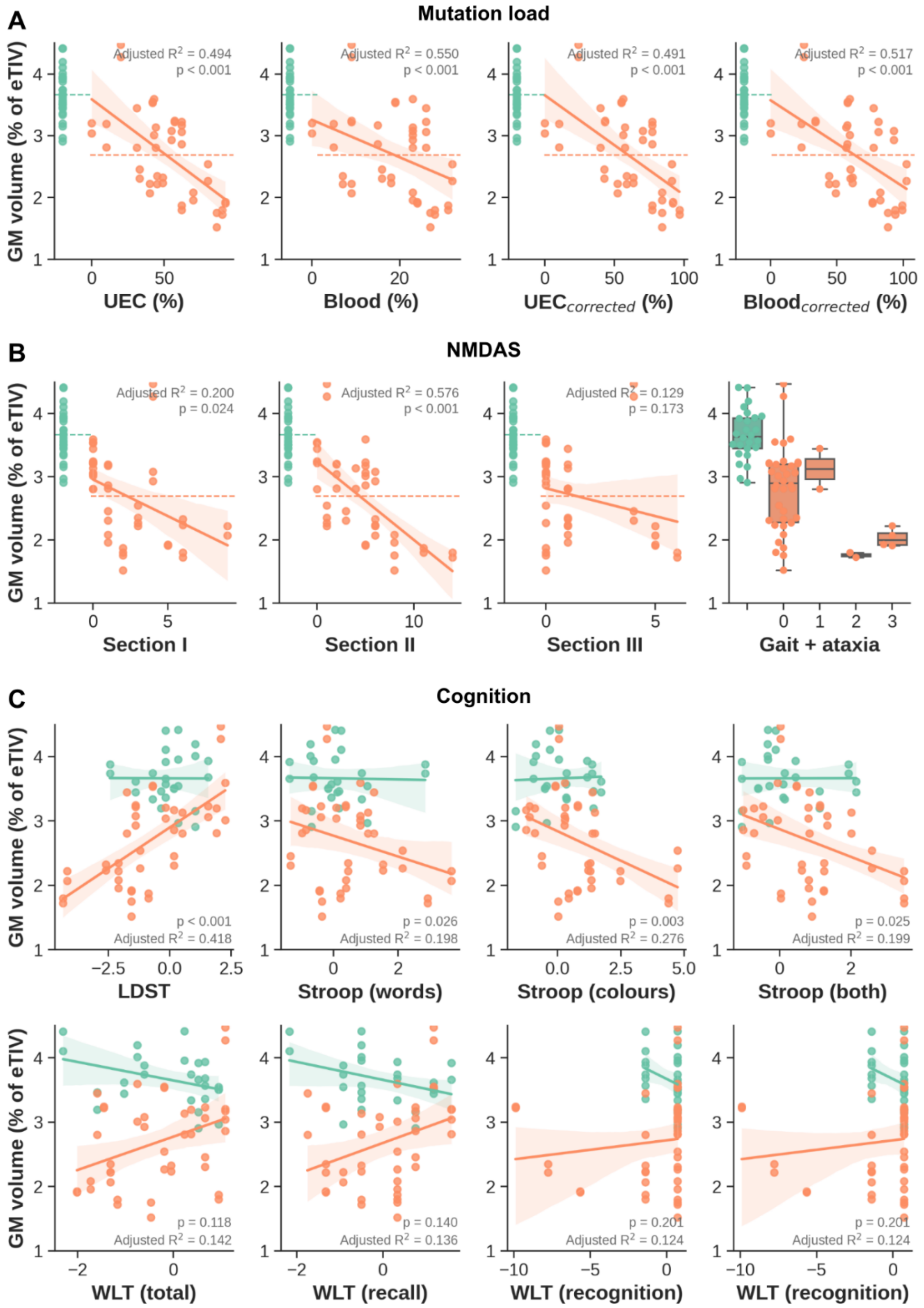


**Supplementary Figure 1 Cerebellar GM volumes.** Correlation between GM volume (y-axis) and (A) uncorrected and corrected mutation load measured in UECs and blood (x-axis), as well as (B) NMDAS subscores per section and across neurological features (x-axis).

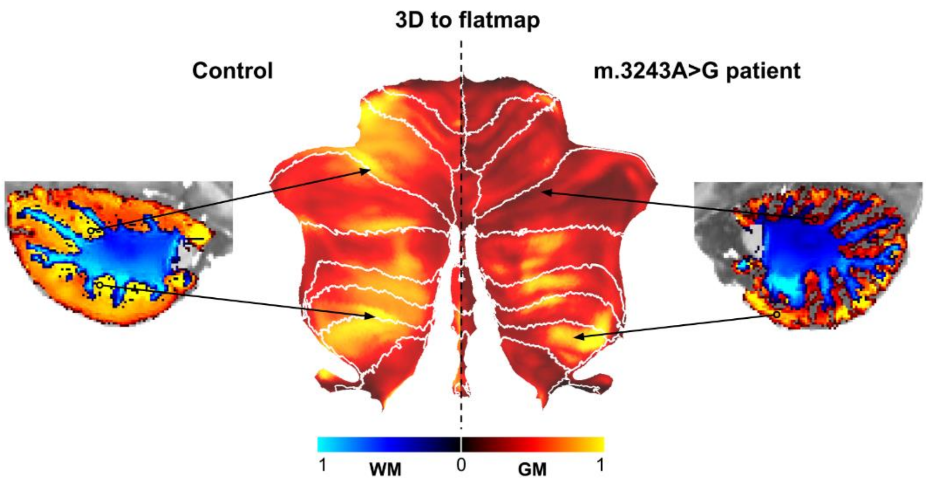


**Supplementary Figure 2 3D to flatmap representation.** GM tissue density maps are mapped from 3D voxel space onto the cerebellar flatmap representation. This example shows the GM tissue density mapping for the same control (left) and m.3243A>G patient (right) as shown in Figure 1 with solid black arrows linking corresponding tissue location in both spaces.

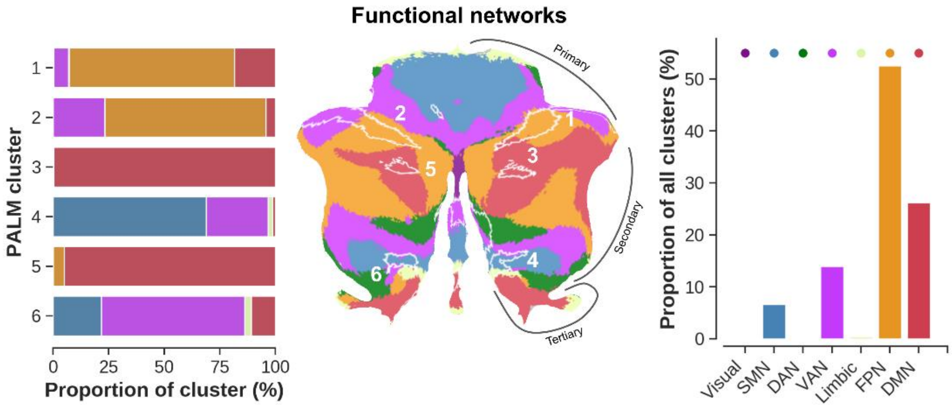


**Supplementary Figure 3 Spatial distribution of the significant clusters with respect to the cerebellar functional networks.** Left to right: stacked bar plot showing statistical (i.e., PALM) clusters (y-axis), ordered from largest at the top (cluster one) to smallest at the bottom (cluster six, in voxels, top x-axis). Here, the width of each individually-colored bar represents the proportional overlap (bottom x-axis) with the respective network. For example, cluster three overlaps entirely with DMN. Middle panel shows a flatmap representation to visualize the localization of each cluster across the cerebellar GM with respect to its lobules. Right panel shows the proportional overlap (y-axis) across all clusters per lobule (x-axis). For example, 50 % of significant voxels fall within FPN.

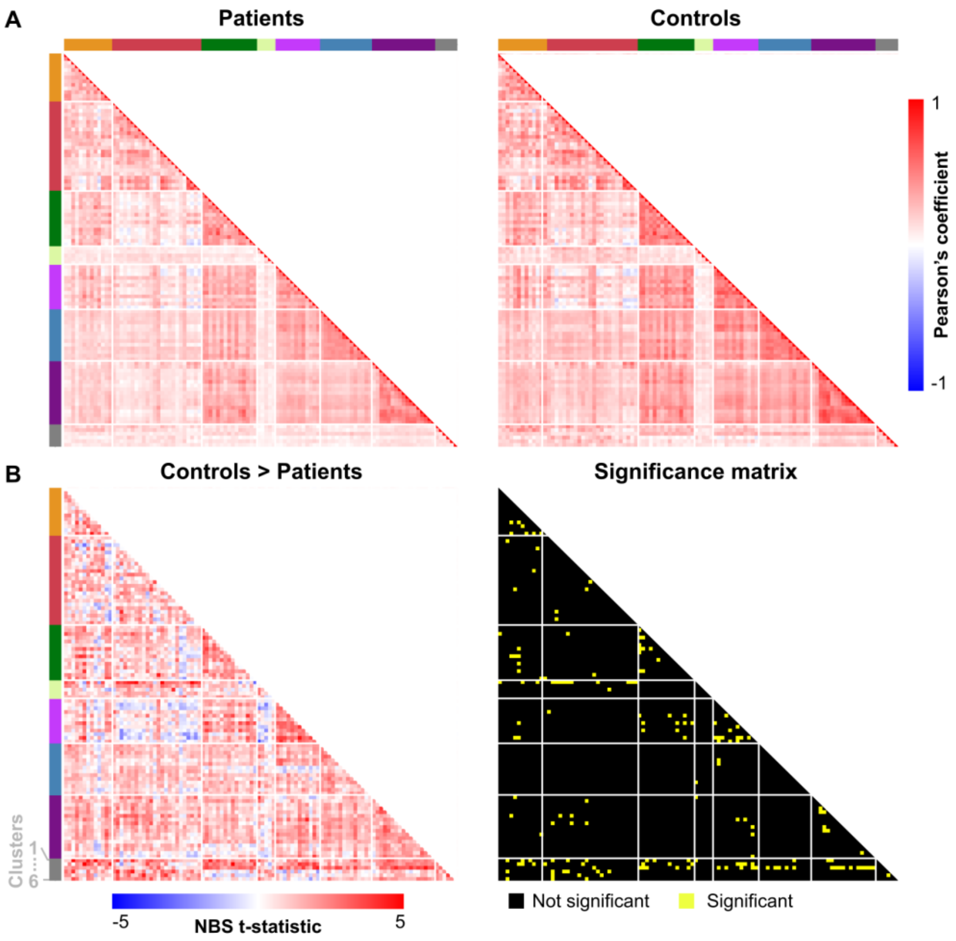


**Supplementary Figure 4 Functional connectivity matrices. (A)** Average Pearson's coefficient connectivity matrices across m.3243A>G patients (left) and controls (right). Each row and column represent a cortical ROI, organized based on functional brain network (top and left colored bars), appended by the statistical clusters in gray. **(B)** Resulting T-statistic (left) and binary significance matrices (right).

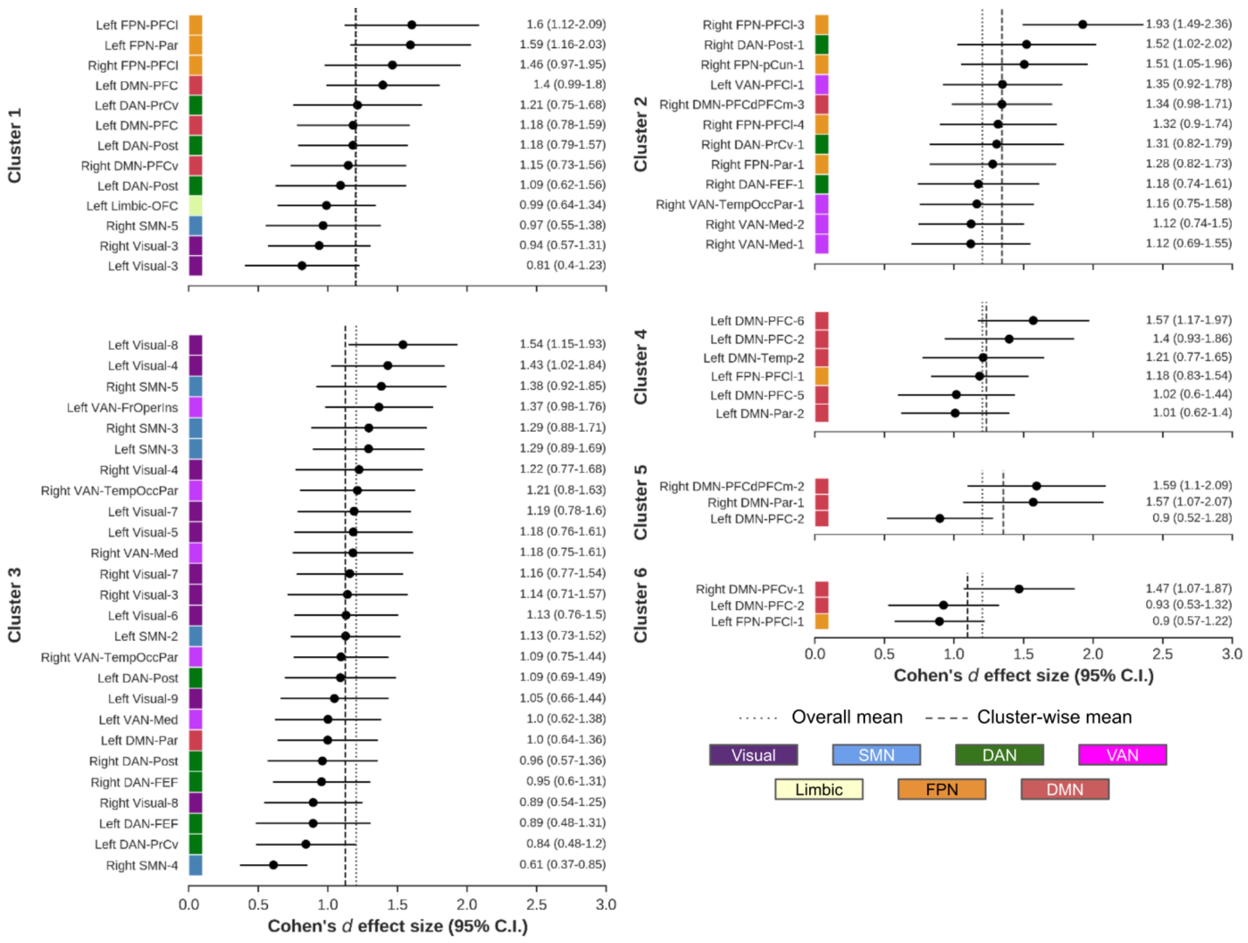


**Supplementary Figure 5 Cluster-wise Cohen's d effect sizes.** Changes in functional connectivity between cerebellar clusters (1-6) and cortical ROIs (y-axes, with color indicating respective large-scale brain network), based on Cohen's d effect size. Only edges significantly impacted in m.3243A>G patients are shown. Vertical dotted and dashed lines indicate overall and cluster-wise mean effect size.

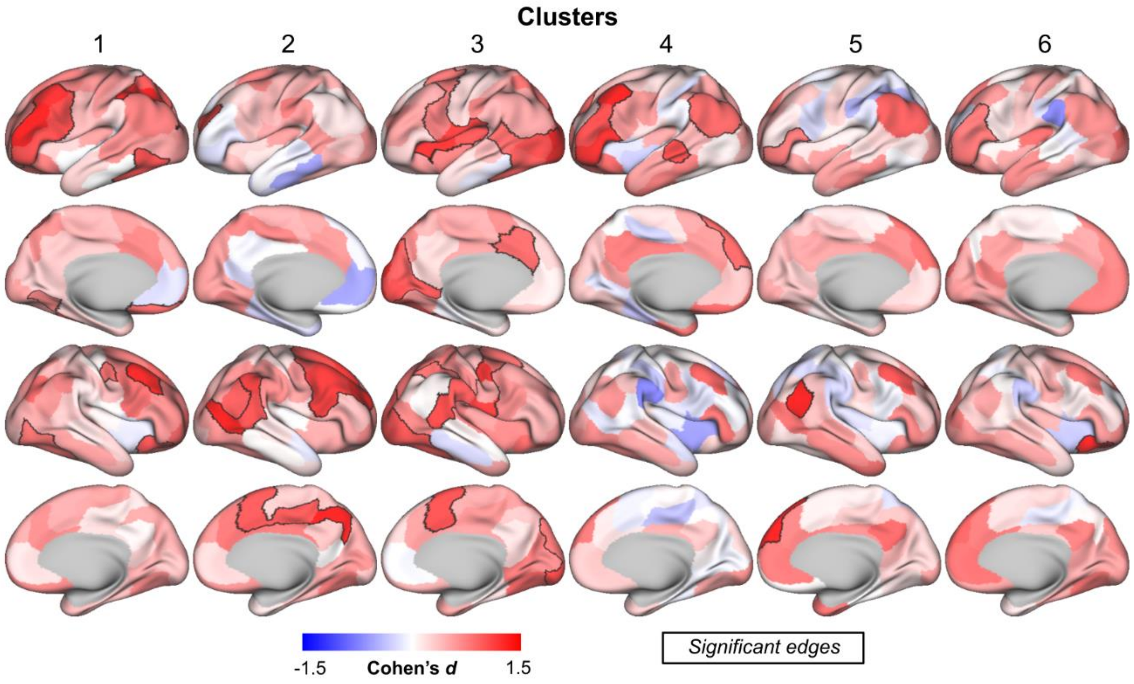


**Supplementary Figure 6 Cluster-wise visualization of functional connectivity changes.** Changes in functional connectivity between each cerebellar clusters (columns) and cortical ROIs, color-coded for corresponding Cohen's d effect size. Edges between clusters and ROIs characterized by significantly reduced connectivity in m.3243A>G patients are delineated using a solid black border.

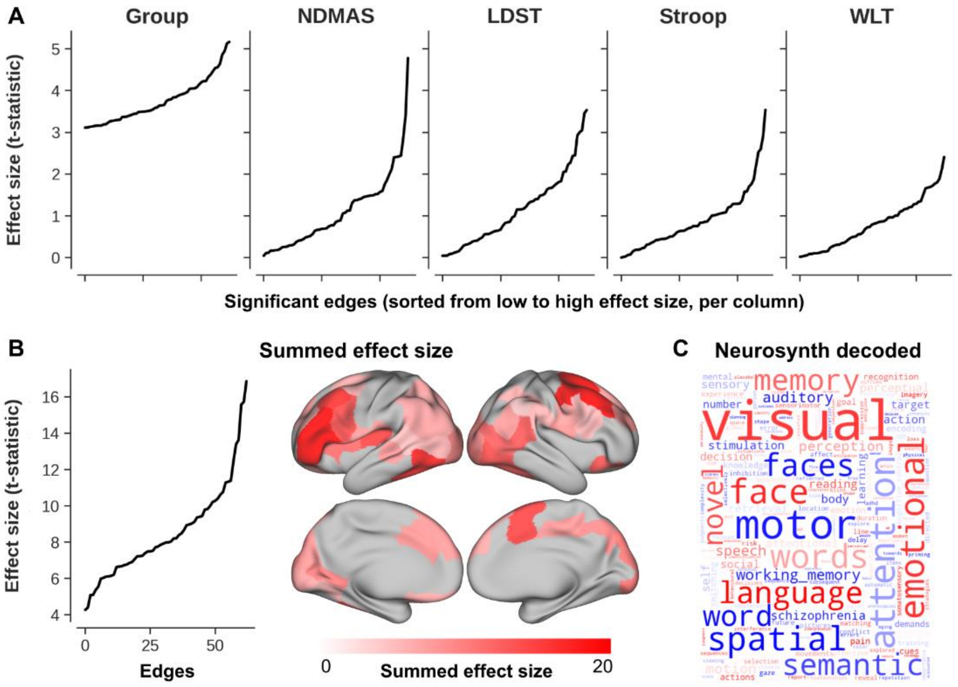


**Supplementary Figure 7 Comparison of effect sizes.** (A) Left to right: line plots with edgewise effect sizes extracted from the comparison between controls and m.3243A>G patients (see Figure 4), and as function of disease severity using NMDAS (see Figure 6) and cognitive performance using LDST, Stroop and WLT (see Figure 7), ordered from smallest to largest maximum effect size. (B) Line plot with edgewise effect sizes summed across data in A and ordered from smallest to largest, as well as displayed onto the cortical surface. (C) Word cloud based on the comparison between the cortical surface map in B and the NeuroSynth database.

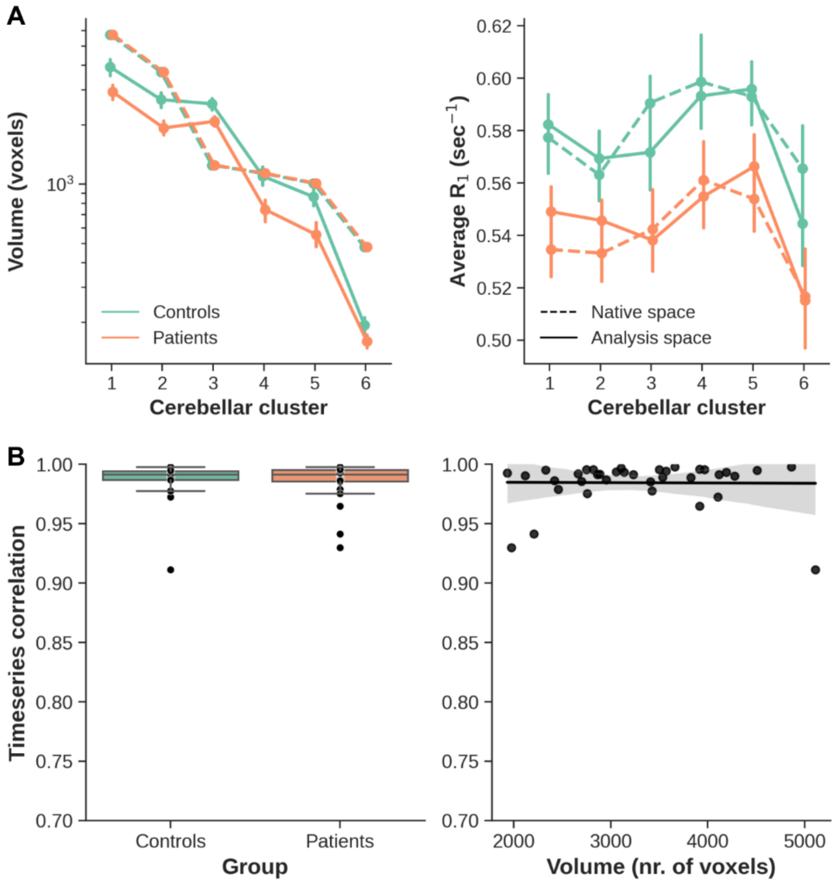


**Supplementary Figure 8 Comparison of volume, $R_1$ and functional connectivity in native and analysis space.** (A) Mapping of the significant cerebellar clusters (x-axis) from analysis (solid lines) back to native (dashed lines) space resulted in significantly different (i.e., controls > patients) cluster volumes (y-axis, left panel). While this is expected because of atrophy (i.e., reduced GM density) in patients for these clusters, the group difference in $R_1$ is expected to be preserved across both spaces. Indeed, the observed pattern of higher $R_1$ values (y-axis, right panel) for controls compared to patients across all clusters (x-axis) was not systematically impacted by the transformation. (B) Correlation between native- and analysis-space data is high (Pearson's r > 0.9), with no apparent difference between patients and controls (x-axis, left panel). Additionally, the correlation coefficient is not dependent on the subject's cluster's volume, and thus, degree of atrophy (x-axis, right panel).
